# A Novel Rotation-Mitigation Technology for Cycling Helmets Tested Across Helmet Types, Impact Locations and Headforms

**DOI:** 10.1101/2025.09.17.676402

**Authors:** Domna-Maria Kaimaki, Higor Alves de Freitas, Archie G. D. Read, Theodore D. M. Dickson, Tony White, Henry C. A. W. Neilson

## Abstract

Head rotation is the leading cause of diffuse brain injuries from cycling accidents, with severe, long-term or even fatal consequences. Here, we present a novel helmet safety technology, the Release Layer System (RLS), designed to enhance conventional helmets and reduce the likelihood of such injuries. RLS is located on the outer side of the helmet and thus gets impacted first. The force of the impact activates a rolling mechanism triggering the release of an outer polycarbonate panel, thereby dispersing and transforming a substantial portion of the incident rotational energy. To evaluate the effectiveness of the technology, we conducted oblique impact tests on three popular helmet types, in conventional and RLS-equipped configurations, at three impact locations. RLS-equipped helmets reduced Peak Angular Velocity (PAV) by 57-66%, averaged across impact locations, compared to their conventional counterparts. This corresponds to a 68-86% reduction in the probability of an AIS2+ brain injury, as estimated by the Brain Injury Criterion. The most notable improvement was observed at the pYrot location (front impacts, mid-sagittal plane) with up to 85% PAV reduction. Testing across headforms further demonstrated the effectiveness of the technology in mitigating head rotation irrespective of variations in evaluation setups. This work introduces a novel mechanism for rotational impact mitigation and provides evidence of its potential benefits compared with conventional helmets. As an outer-layer approach, RLS may offer an alternative pathway for managing rotational kinematics in future helmet designs.

## 1 Introduction

Within the last decade, cycling has experienced a notable rise in popularity with the global number of bicycles estimated to be over a billion ^1,2^. As cycling activity increases, so does the number of associated incidents. In Europe, cycling is the only mode of transport that has shown almost no decline in fatalities over the past decade, while also making up 24% of all serious injuries in 2020^3^. In response, a multidisciplinary body of research has emerged to better understand and mitigate risks within an increasingly complex and dynamic road environment. The interaction between Vulnerable Road Users (VRUs) and motorised vehicles has been further investigated using video data for accident reconstructions ^4,5^, while comprehensive databases - integrating police reports, clinical records, and insurance claims - have been developed to support injury analysis^6,7^. At the same time, advances in sensor technology and computer vision methods are being used to suggest infrastructure updates and explore real-time accident prevention strategies ^8^, and policy recommendations are increasingly being derived from case studies in countries such as the Netherlands, Denmark and Germany ^9–11^.

While systemic changes to infrastructure and policy are crucial, interventions at the individual cyclist level offer more immediate and actionable safety benefits. Cyclist head injuries are consistently identified as among the most common injury types and as the leading contributor to severe and fatal outcomes in cycling crashes, making helmets the primary focus of cyclist protective equipment ^12–14^. Biomechanical studies have shown that wearing a helmet is indeed effective in reducing the transmission of impact forces to the head and, in doing so, lowering the risk of Traumatic Brain Injury (TBI) compared with unhelmeted impacts ^15,16^. This protective ability of helmets is assessed via standardised linear impact tests, as defined by certification protocols, such as EN 1078 for Europe, CPSC 1203 for the US, AS/NZS 2063 for Australia and New Zealand, and JIS T 8134 for Japan ^17–20^. However, linear impact testing only replicates focal head injuries, such as skull fractures, neglecting head rotation and its effect to brain injury, which has long been established ^21^. Advances in research connecting medical imaging and clinical outcomes to brain modelling and biomechanics, have shown that head rotation is responsible for concussions, certain types of haematomas, and diffuse axonal injuries (DAIs) ^22^. Such pathologies may arise from mild Traumatic Brain Injuries (mTBIs), but they have been linked to long-lasting impairments in sleep quality, motor function, memory, as well as more severe neurological damage and even death ^23^. As a result, efforts to adapt helmet certification standards to reflect real-world oblique impacts, inducing both linear and rotational head kinematics, are currently underway. In parallel with evolving certification requirements, helmet manufacturers have also introduced various new materials, design refinements, and additional technologies into helmets to complement their linear impact protection by mitigating rotational energy transfer^24,25^. It is within this context that we introduce the Release Layer System (RLS) technology for cycling helmets.

### 1.1 State of the art & the Release Layer System (RLS)

Conventional helmets typically consist of at least two material layers: (i) an outer shell - commonly a thermoformed sheet of polycarbonate - that sustains the impact, distributes forces over a larger area, and absorbs energy through deformation or fracture, and (ii) a liner, often made of expanded polystyrene (EPS) that absorbs and dissipates energy through compression, providing cushioning to minimise head injuries. Over the past two decades, conventional helmets have been supplemented with additional materials and structures aimed at mitigating rotational head kinematics during an oblique impact to address the leading cause of mTBIs. Technologies such as MIPS, SPIN, WaveCel, Koroyd, KinetiCore, and Rheon have been integrated into helmet interiors, reflecting an evolution in helmet design intended to offer a meaningful advancement in the helmet’s effectiveness in addressing rotational loading ^26–31^.

Here, we present a different approach to rotation mitigation: an outer technology, called the Release Layer System (RLS). The RLS technology consists of an array of polycarbonate spheres of 2.0 ± 0.1 mm in diameter, attached onto a flexible membrane and encapsulated between the outer polycarbonate shell of a conventional EPS helmet (‘B-surface’) and an additional polycarbonate shell (‘A-surface’) split into panels. The polycarbonate spheres are held in place by adhesives on the top and bottom, while the assembly of membrane and spheres is secured onto a helmet by a combination of bonding with the B-surface using an adhesive and mechanical interlocking with the A-surface using fasteners (Figure 1A). During an impact, the oblique force on the A-surface panel severs the mechanical fasteners and the adhesive connections of the spheres in the impacted area allowing the spheres to roll freely, while the panel releases from the B-surface. This mech-anism operates on a principle analogous to that of ball bearings, enabling relative rotational motion between outer helmet layers through a rolling interface. The rolling of the spheres disperses and transforms a portion of the energy of the impact, thereby protecting the head from rotational acceleration (Figure 1B), and is complemented by the EPS liner, which absorbs part of the remaining energy through compression ^32^.

**Fig. 1.**
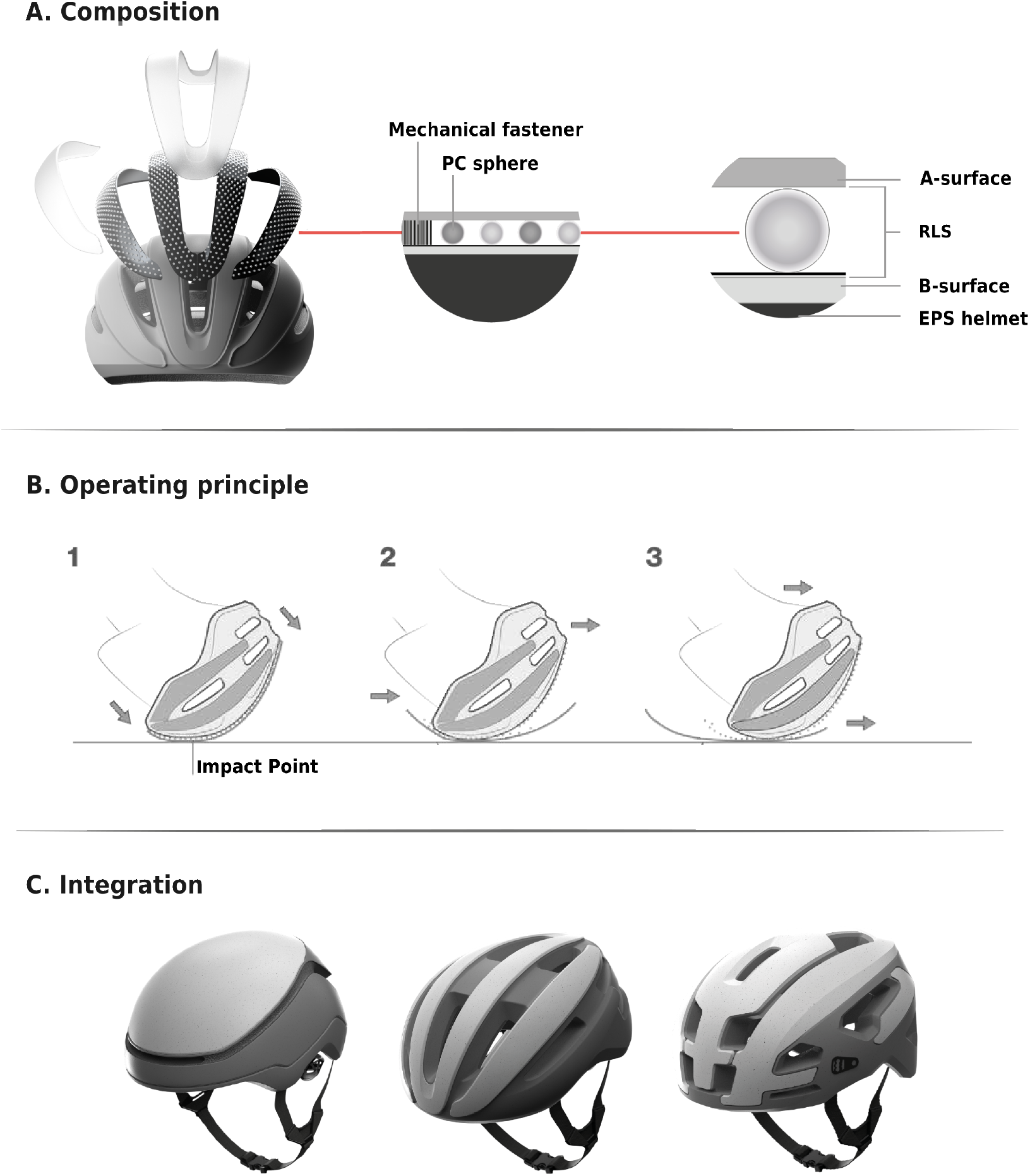
The Release Layer System (RLS) in cycling helmets. A. A deconstructed RLS-equipped helmet presenting the composition of the technology, B. The operating principle of the RLS technology during an impact, C. Integration of the RLS technology onto the different helmet types tested in this study; the urban, road and mountain bike helmets are displayed from left to right.

RLS can be integrated into a variety of helmets; in this work it has been implemented in urban, road and mountain-bike (MTB) helmets (helmet types making up 83% of all helmets rated by Virginia Tech Helmet Ratings ^33^). In urban helmets, RLS is configured as a single membrane with spheres between surfaces A and B. In road and MTB helmets, RLS is divided into multiple sections of the same composition, to accommodate helmet features, such as vents, that are crucial for helmet comfort. This is mirrored in the A-surface, which is likewise partitioned into multiple panels. In this publication, we present helmets with one, three and four panels for urban, road and MTB helmets, respectively (Figure 1C), and experimentally assess the effectiveness of the RLS technology in mitigating head rotation by performing oblique impact testing of conventional and RLS-equipped cycling helmets of all three types. The study is structured into two parts: (i) an evaluation of RLS effectiveness across three cycling helmet types and three impact locations using a Hybrid III (HIII) headform, and (ii) an assessment of the sensitivity of RLS effectiveness, within a single helmet type, to headform selection through testing with the EN 17950 headform.

## 2 Materials and Methods

### 2.1 Helmets

All urban, road and MTB helmets - in their conventional and RLS-equipped configurations - used in this study were manufactured by HEXR Ltd (London, UK) or its partners. The conventional helmets comprised of EPS (nominal density of 70 g*/*l) and a polycarbonate shell, similarly to the majority of cycling helmets on the market. The polycarbonate shell was formed to have cavities that could accommodate the RLS technology. The RLS-equipped helmets had the same composition and geometry as the conventional helmets with the addition of the RLS technology and the A-Surface on the outer side (Figure 1A, C, Table S1 in Supplementary Information).

### 2.2 Headforms & instrumentation

The majority of the experiments in this publication were conducted with the 50th percentile male HIII headform (58 cm head circumference, Humanetics Innovative Solutions Inc, Michigan, United States) that has been widely used for the evaluation of helmets ^27,28,34^. These experiments were conducted at the iCUBE laboratory of the University of Strasbourg (UNISTRA), where the HIII headform is equipped with a triaxial linear accelerometer (PCB Piezotronics, New York, United States) and angular rate sensors (ATA Sensors, New Mexico, United States) measuring in the SAE J211 orientation. The accelerometers are located at the center of gravity of the headform, have an amplitude range of ±500 g and a sampling rate of 25.6 kHz by channel, while the an-gular rate sensors have a measurement capacity of ±8000 deg*/*s, a bandwidth of 1 kHz and a sampling rate of 25.6 kHz. After each impact, the HIII headform was cleaned using a dry fiber cloth to remove any debris or particles transferred from the tested helmet.

Additional testing was conducted using a second headform, the EN 17950 (57 cm head circumference, Humanetics Innovative Solutions Inc, Michigan, United States). This headform was developed under the instruction of the European Committee for Standardization Working Group 11 (CEN/TC158/WG11) with the aim to be a more biofidelic model of a human head, both in terms of its moments of inertia (MoIs) and its coefficient of friction (CoF), addressing criticisms faced by the other available headforms ^35–37^. Testing with the EN 17950 headform was conducted at the HEXR laboratory due to the limited availability of the headform, compatible instrumentation, and related equipment (eg. CoF measurement setup) in other laboratories at the time of the study. Prior to testing, the EN 17950 headform was cleaned following instructions by the manufacturer and its CoF was quantified against a structured polyester strap complying with EN 12195-2, as specified in the BS EN 17950:2024 standard ^38^. The CoF was measured before and after testing (all tests were completed in one day) and the mean of each test sequence was found to be within the acceptable range (0.30 ± 0.03 N*/*m, Table S2 in Supplementary Information); during testing the headform was handled with gloves to minimise contamination. The EN 17950 headform is instrumented with a 6DX Pro-A system (DTS, California, United States) including a triaxial accelerometer (amplitude range of ±500 g and bandwidth of 5 kHz) and angular rate sensor (amplitude range of ±8000 deg*/*s and bandwidth of 2 kHz) with a sampling rate of 20 kHz. As the instrumentation is not located at the headform’s center of gravity, the measured linear acceleration data was transformed to that point using rigid body kinematic equations provided by the manufacturer.

### 2.3 Impact testing methodology

Oblique impact testing was conducted using a drop tower (AD Engineering at UNISTRA and custom-made at the HEXR laboratory) comprising of a trolley to hold the helmet-headform assembly, stainless steel rails to guide the trolley with the testing assembly to the appropriate height, and a 45 ± 0.5° stainless steel anvil (diameter of 130 ± 3 mm). The oblique anvil has the same impact area and material specifications as the flat anvil specified in the EN 1078 standard ^17^ but is angled at 45° following the rationale explained in CEN/TR 18249^39^. The anvil surface was covered with P80 abrasive paper to simulate the road surface; the paper was replaced once it was visibly damaged. In order to assess the variability of the results, we conducted 4 repeats at each of the 3 impact locations (pXrot, pYrot and pZrot) for each helmet type (urban, road, MTB) and configuration (conventional and RLS-equipped). This resulted in *n* = 72 impacts with the HIII headform at UNISTRA. Testing at the HEXR laboratory was conducted with the same number of repeats but using only the urban helmet resulting in *n* = 24 impacts with the EN 17950 headform (Table S1 in Supplementary Information). Each helmet was tested only once to assess its effectiveness against a primary impact to the head. MTB helmets were tested without visors ^31,40^.

Prior to testing, the helmets were stored at a room temperature of 20 ± 2°C and a relative humidity of 50 ± 20% for a period of at least 4 hours; conditions that were maintained throughout the duration of the experiments. The headform was first aligned with the helmet using a nose gauge (indicating a normal distance of 63 mm between the tip of the nose and the helmet brim, Figure 2A). The impact speed for all testing was set to 6.5 m*/*s and tests were conducted on three locations chosen to represent the most common head impact locations in cycling accidents ^22,41,42^. The headform-helmet system was positioned in the drop tower according to the impact location required for testing. For pXrot and pYrot impacts, the central vertical axis of the headform (z-axis) was aligned to the vertical with a tolerance of ±1°, while the assembly (xy-plane) was placed at 0 ± 1° around the z-axis from the front. The normal vector of the impact surface was parallel to the yz-plane and xy-plane, for pXrot and pYrot, respectively with a tolerance of ±1°. For pZrot impacts, the assembly was inclined by 65 ± 1° around the y-axis from the base of the headform with the horizontal plane (the yz-plane was placed at 25 ± 1° from the horizontal, Figure 2B). Inclinometers (High Precision Digital Level TLL-90S, Jingyan Instruments, China) were used in both laboratories to ensure alignment. This methodology follows closely the oblique testing protocol implemented by Folksam with an increase of the impact speed to 6.5 ± 0.1 m*/*s to align with the higher impact speed suggested in the provisional EN 1078:2025 standard (^34^)

**Fig. 2.**
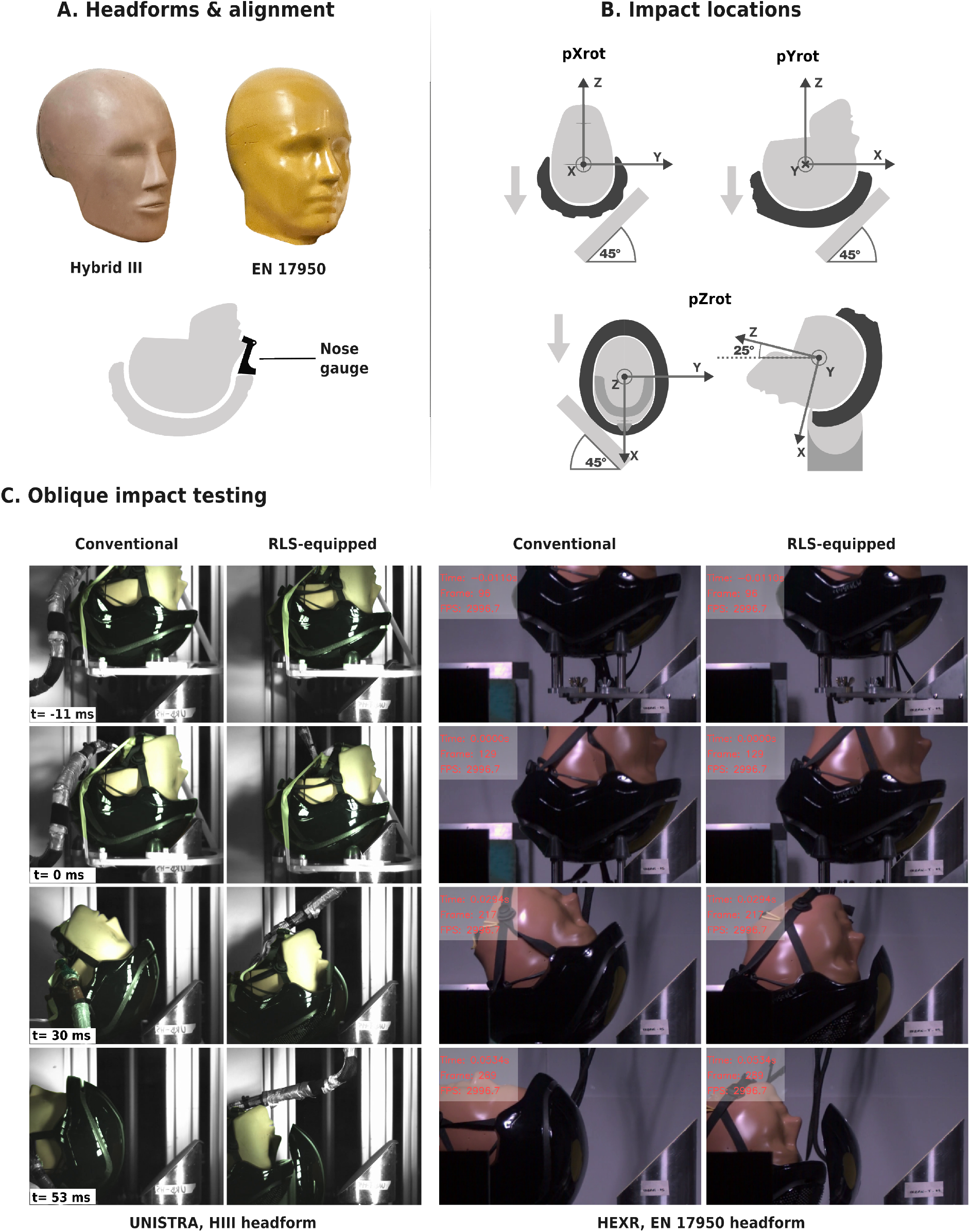
The testing methodology of this study. A. The two headforms (HIII & EN 17950) that were used for impact testing and the nose gauge of 63 mm that was used for alignment of the helmet-headform assembly. B. The alignment of the headform-helmet assembly on the anvil for each impact location (pXrot, pYrot and pZrot) used for oblique testing. C. Snapshots of the high speed videos during oblique impact testing at UNISTRA and the HEXR laboratory for a visual comparison of rotation on conventional and RLS-equipped helmets.

### 2.4 Data acquisition and analysis

Once alignment was ensured, the headform-helmet assembly was secured in place to prevent it from moving as the trolley is raised (using tape at UNISTRA and a pneumatic actuator at HEXR). Subsequently, the instrumentation was armed and the trolley was raised to the required height as determined by a speed-height calibration protocol. The trolley was then released initiating a free fall of the assembly, the impact was recorded and the resulting impact velocity was measured using a light gate (BeeSpi V Light Gate, Vitta Education, United Kingdom). Additionally, a high speed video (CR3000x2, Optronis GmbH, Germany at UNISTRA and Chronos 2.1 HD High Speed Camera, Kron Technologies, Canada at HEXR) was recorded to visualise the impact and to further assess the alignment of the headform-helmet assembly on the drop tower post-impact (Figure 2C). An AD Engineering data acquisition system and Sliceware were used at UNISTRA and the HEXR laboratory, respectively to collect and preprocess the kinematic data. The raw data from both laboratories was filtered with a CFC 1000 for linear acceleration and a CFC 180 for angular velocity, in accordance with ISO 6487^43^.

The results are presented with a focus on peak values of key kinematic metrics: peak linear acceleration (PLA), peak angular velocity (PAV) and peak angular acceleration (PAA), where the angular acceleration is calculated by numerical differentiation from the angular velocity data (forward difference with a step of 1 over sampling rate and no smoothing). To quantify the variability of these metrics, descriptive statistics such as standard deviation and coefficient of variation (CV) were calculated. The effectiveness of the RLS technology was quantified as the mean percentage reduction in such metrics in comparison to conventional helmets tested with the same headform:

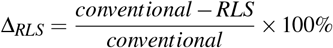

Inferential comparisons between RLS-equipped and conventional helmets were conducted using statistical tests selected based on the underlying data characteristics. Specifically, the Shapiro-Wilk test and Levene’s test were used to assess the assumptions of normality and homoscedasticity, respectively. Based on the outcome of these tests, Welch’s ANOVA was used when the data was normally distributed as it provides improved control of Type I error, whereas the Mann-Whitney U test was employed for non-normally distributed data. In the second part of the study, the effectiveness of RLS was quantified for two headforms and a permutation test was employed to test the hypothesis that HIII and EN 17950 headforms yield exchangeable results of RLS effectiveness. The test statistic for the permutation test is defined as the difference in mean percentage reduction,

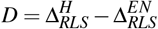

Given the small sample size and the nonlinear nature of the estimator, hypothesis testing was performed using an exact permutation test (unpaired) with headform labels assumed to be exchangeable within each configuration (conventional and RLS-equipped). All possible permutations were enumerated (4900 exact permutations per metric), and a two-sided p-value was calculated as the proportion of permutations for which |*D*| was greater than or equal to the observed value. A permutation-based 95% Confidence Interval (CI) for *D* was also computed and reported. Statistical significance for all statistical tests was set at *α* = 0.05. All statistical analyses were conducted using custom Python code (v3.10.13) including the following libraries: pandas, numpy, scipy, matplotlib, statsmodels and pingouin. Beyond the evaluation of peak kinematics, the data was also used to assess the potential of the RLS technology to mitigate brain injuries. For this purpose, we employed the Brain Injury Criterion (BrIC) incorporating direction-specific thresholds for critical angular velocities. These thresholds were based on values derived from the cumulative strain damage measure (CSDM), maximum principal strain (MPS), and their average, as reported by Takhounts et al ^44^. The resulting BrIC values were used to estimate the probability of sustained Abbreviated Injury Scale 2 or higher (AIS2+) injuries (Equations 1 and 2 in Supplementary Information).

## 3 Results

### 3.1 Average RLS effectiveness by helmet type (HIII headform)

The performance of the conventional helmets was examined first as they were used to establish the baseline throughout this study. The average PLA across all conventional helmets was 116-127g, the average PAV was 33-34 rad/s, while the average PAA was 6.7-7.3 krad*/*s^2^. On the other hand, the average PLA across all RLS-equipped helmets was 109-116g (Δ_*RLS*_ of 4.5-9.7%). This reduction is not significant for urban and MTB helmets (*U* = 97.0, *p* = 0.157 and *U* = 92.0, *p* = 0.260, respectively), but it is significant for road helmets (*U* = 129.0, *p* = 0.001). In terms of rotational kinematics, the average PAV across all RLS-equipped helmets was 11-15 rad/s (Δ_*RLS*_ of 57-66%), while the average PAA was 2.9-4.1 krad*/*s^2^ (Δ_*RLS*_ of 44-56%).

For each helmet type, peak kinematic metrics were first averaged across the three tested impact locations to provide a location-agnostic measure of overall RLS effectiveness, enabling direct comparison between helmet types. Examination of the results (Table 1) shows that the largest reduction in rotational kinematics relative to conventional helmets was observed for the road helmet type (66% reduction in PAV and 56% in PAA), followed by the MTB helmet type (58% reduction in PAV and 47% in PAA) and finally the urban helmet type (57% reduction in PAV and 44% in PAA). This trend can also be visualised in Figure 3, where the empty diamonds show the mean across all three impact locations for each helmet type and statistical differences are indicated with an asterisk. The decrease in rotational kinematics observed for RLS-equipped helmets of all three helmet types corresponds to a reduction in the average risk of sustaining an AIS2+ injury across impact locations of 68-86%. In simple terms, based on headform-measured kinematics and associated injury criteria, these results indicate that, under the tested impact conditions a cyclist wearing the conventional urban, road or MTB helmets tested in this study is 68%, 64% or 64% likely to suffer a mTBI of AIS2+, such as a concussion or even loss of consciousness of up to 1 hour, if involved in an accident ^45^. The probability of a similar injury be-ing sustained if the cyclist wears the corresponding RLS-equipped helmet is significantly reduced to 22%, 9% and 13% for the urban, road, or MTB helmet, respectively.

**Table 1.** Summary of data from oblique testing with a HIII headform. Mean values and standard deviations were calculated by averaging across three locations (n=12 datapoints) for each helmet type and configuration (control, RLS). All PAV, PAA, BrIC and p(AIS2+) comparisons between conventional and RLS-equipped helmets were statistically significant with *U* = 126.0 − 144.0 and *p* ≤ 0.002. In contrast, PLA comparisons between configurations were only statistically significant for the road helmet type (Mann-Whitney U test results in bold).

| Metric | Urban |  | Road |  | MTB |  |
| --- | --- | --- | --- | --- | --- | --- |
|  | Control | RLS | Control | RLS | Control | RLS |
| PLA (g) | 121.8 ± 7.1 | 116.3 ± 7.1 | 127.3 ± 7.0 | 115.0 ± 6.2 | 115.5 ± 12.9 | 109.3 ± 12.7 |
| | $U = 97.0, p = 0.157$ | | **$U = 129.0, p = 0.001$** | | $U = 92.0, p = 0.260$ | |
| PAV (rad/s) | 34.2 ± 2.8 | 14.7 ± 7.9 | 33.2 ± 1.6 | 11.3 ± 3.1 | 32.8 ± 0.8 | 13.7 ± 3.7 |
| PAA (krad/s<sup>2</sup>) | 7.32 ± 1.67 | 4.12 ± 2.12 | 6.69 ± 0.49 | 2.92 ± 1.24 | 6.81 ± 1.10 | 3.60 ± 1.05 |
| BrIC | 0.65 ± 0.13 | 0.32 ± 0.18 | 0.62 ± 0.11 | 0.24 ± 0.09 | 0.61 ± 0.10 | 0.28 ± 0.09 |
| p(AIS2+) | 0.68 ± 0.18 | 0.22 ± 0.21 | 0.64 ± 0.18 | 0.09 ± 0.09 | 0.64 ± 0.16 | 0.13 ± 0.11 |

**Fig. 3.**
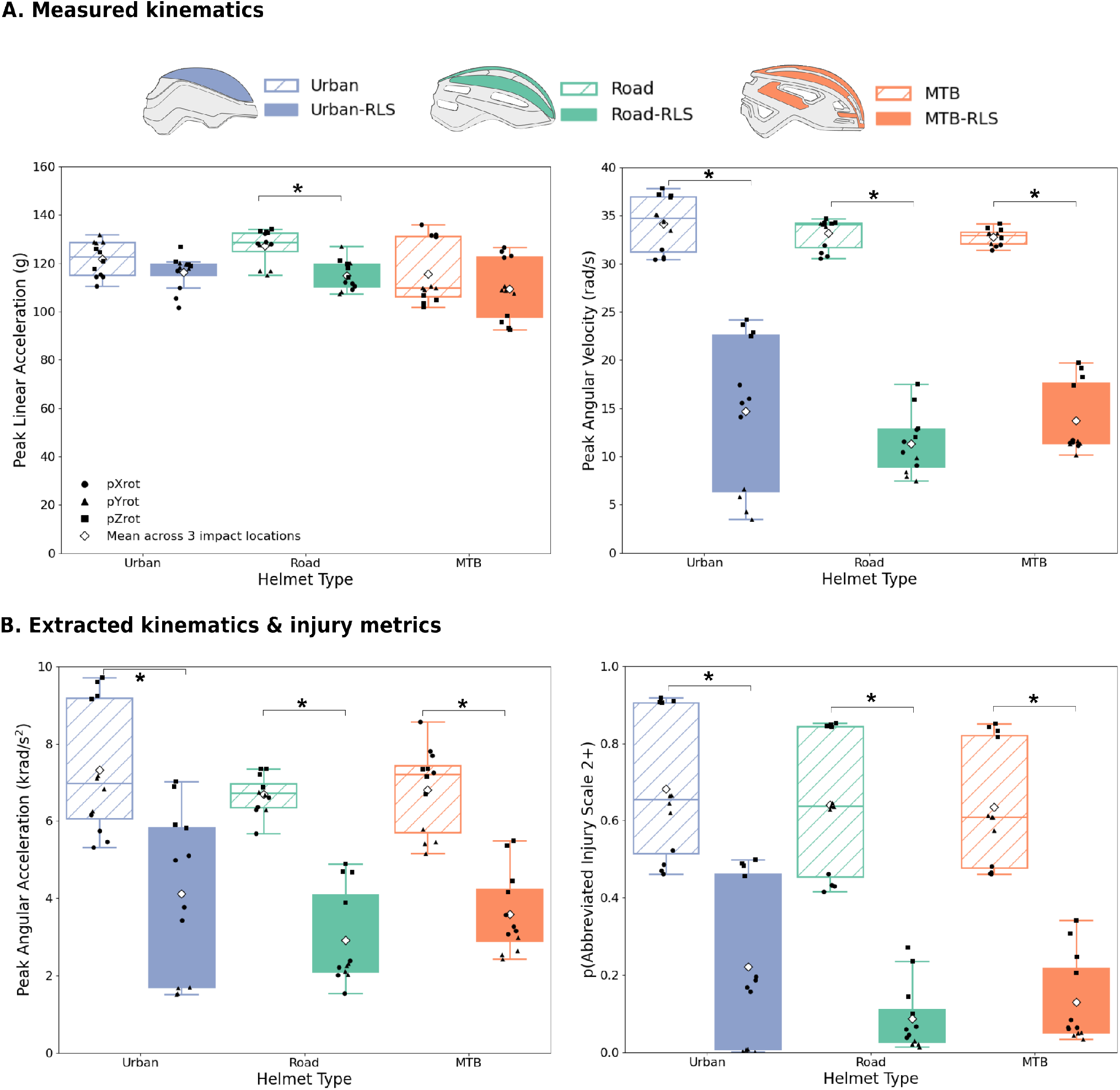
Data for conventional (hatched) and RLS-equipped (solid) helmets, grouped by helmet type (colour-coded), obtained using a HIII headform and averaged across the three tested impact locations. Measured peak kinematics (PLA, PAV) are shown in A, and calculated peak kinematics and injury metrics (PAA, p(AIS2+)) are shown in B. Empty diamonds indicate mean values across impact locations, while black circles, triangles, and squares represent results for the pXrot, pYrot, and pZrot impact locations, respectively. Asterisks (*) denote statistically significant differences (*p <* 0.05).

However, assessing the effectiveness of the RLS technology for each helmet type by solely considering the average values across impact locations is not sufficient. As illustrated in Figure 3, the results vary with impact location (black circles, triangles and squares representing pXrot, pYrot and pZrot results, respectively). The urban helmet shows the highest variability in all rotational kinematics and injury metrics between locations, while the road and MTB helmets have a narrower spread. The implications of this variability for Δ_*RLS*_ are explored in the following section.

### 3.2 RLS effectiveness by impact location for three helmet types (HIII headform)

The data collected for conventional and RLS-equipped helmets was analysed per impact location with pXrot simulating side impacts on the frontal plane, while pYrot and pZrot are simulating front impacts on the mid-sagittal and parasagittal planes, respectively (Figure 2B). All comparisons of peak rotational kinematics and extracted injury metrics between conventional and RLS-equipped helmets were statistically significant, irrespective of the helmet type (*U* = 16.0, *p* = 0.029 in Table 2, indicated by the asterisk in Figure 4).

**Table 2.** Kinematic and injury metric data from oblique testing with a HIII headform. Mean values and standard deviations were calculated by impact location for each helmet type and configuration (n=4 datapoints). All PAV, PAA, BrIC and p(AIS2+) comparisons between conventional and RLS-equipped helmets were statistically significant with *U* = 16.0 and *p* = 0.029. In contrast, PLA comparisons between configurations were only statistically significant in certain impact locations for certain helmet types and thus, the Mann-Whitney U test results are included in the table. Statistically significant results are indicated in bold.

| Metric | Urban |  |  |  |  |  |
| --- | --- | --- | --- | --- | --- | --- |
|  | pXrot |  | pYrot |  | pZrot |  |
|  | Control | RLS | Control | RLS | Control | RLS |
| PLA (g) | 113.7±2.1<br>$U = 12.0, p = 0.343$ | 108.5±6.6 | 129.5±1.5<br>$U = 16.0, p = 0.029$ | 119.0±1.1 | 122.2±3.7<br>$U = 9.0, p = 0.886$ | 121.5±3.6 |
| PAV (rad/s) | 30.8±0.5 | 15.8±1.4 | 34.5±0.8 | 5.1±1.4 | 37.3±0.4 | 23.3±0.8 |
| PAA (krad/s<sup>2</sup>) | 5.67±0.37 | 4.33±0.85 | 6.85±0.42 | 1.61±0.10 | 9.43±0.27 | 6.41±0.64 |
| BrIC | 0.52±0.01 | 0.34±0.01 | 0.61±0.01 | 0.09±0.02 | 0.82±0.01 | 0.52±0.01 |
| p(AIS2+) | 0.49±0.03 | 0.18±0.02 | 0.65±0.02 | 0.01±0.00 | 0.91±0.01 | 0.48±0.02 |
| Metric | Road |  |  |  |  |  |
|  | pXrot |  | pYrot |  | pZrot |  |
|  | Control | RLS | Control | RLS | Control | RLS |
| PLA (g) | 129.4±2.0<br>$U = 16.0, p = 0.029$ | 110.9 ±1.3 | 119.2±5.7<br>$U = 10.0, p = 0.686$ | 115.7 ±9.6 | 133.2±0.7<br>$U = 16.0, p = 0.029$ | 118.3 ±2.9 |
| PAV (rad/s) | 31.1±0.6 | 11.0 ±1.6 | 34.1 ±0.2 | 8.4 ±1.0 | 34.3±0.2 | 14.6±2.6 |
| PAA (krad/s<sup>2</sup>) | 6.24±0.39 | 2.04±0.37 | 6.65±0.25 | 2.18±0.13 | 7.20±0.22 | 4.54±0.45 |
| BrIC | 0.49±0.01 | 0.21±0.02 | 0.61±0.00 | 0.16±0.02 | 0.75±0.00 | 0.34±0.06 |
| p(AIS2+) | 0.44±0.02 | 0.05±0.01 | 0.64±0.01 | 0.02±0.01 | 0.85±0.00 | 0.19±0.08 |
| Metric | MTB |  |  |  |  |  |
|  | pXrot |  | pYrot |  | pZrot |  |
|  | Control | RLS | Control | RLS | Control | RLS |
| PLA (g) | 132.6±2.3<br>$U = 16.0, p = 0.029$ | 124.3±1.9 | 109.8±0.5<br>$U = 12.0, p = 0.343$ | 108.9 ±1.3 | 104.2 ±2.0<br>$U = 16.0, p = 0.029$ | 94.9±2.7 |
| PAV (rad/s) | 32.0±0.5 | 11.4±0.2 | 32.9±0.5 | 11.1±0.6 | 33.5±0.6 | 18.6±1.0 |
| PAA (krad/s<sup>2</sup>) | 7.83±0.54 | 3.27 ±0.22 | 5.46±0.26 | 2.65±0.24 | 7.13±0.30 | 4.87±0.67 |
| BrIC | 0.51±0.00 | 0.24±0.01 | 0.58±0.01 | 0.20±0.01 | 0.74±0.01 | 0.40±0.04 |
| p(AIS2+) | 0.47±0.01 | 0.07±0.01 | 0.60±0.02 | 0.04±0.01 | 0.84±0.01 | 0.28±0.06 |

**Fig. 4.**
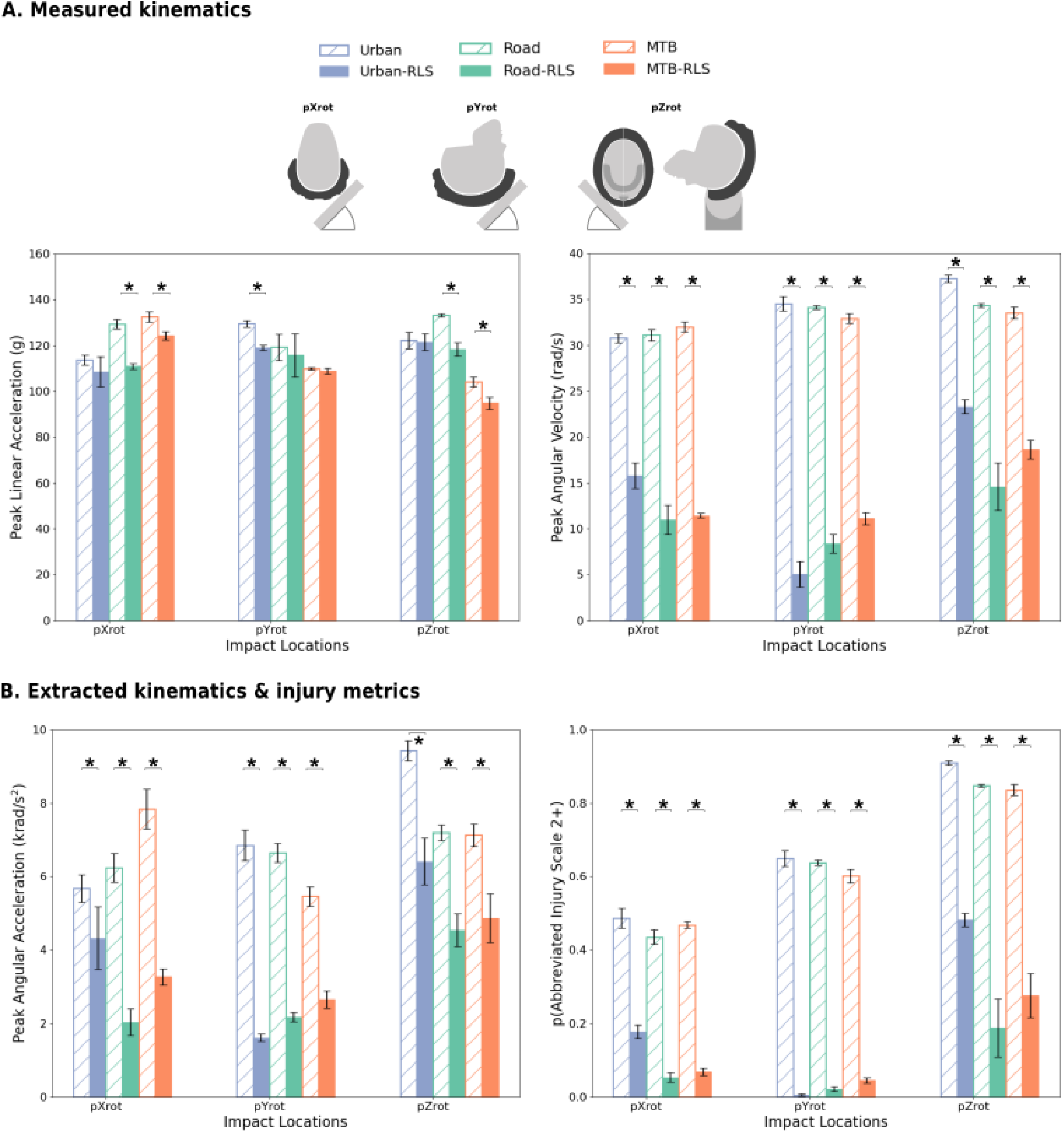
Data for conventional (hatched) and RLS-equipped (solid) helmets by impact location and helmet type (colour-coded), obtained using a HIII headform. Measured peak kinematics (PLA, PAV) are shown in A, and calculated peak kinematics and injury metrics (PAA, p(AIS2+)) are shown in B. Asterisks (*) denote statistically significant differences (*p <* 0.05).

The largest Δ_*RLS*_ in PAV was observed at the pYrot location (85%, 75% and 66% for the urban, road and MTB helmets, respectively), followed by the pXrot location (49%, 65% and 64%) and finally the pZrot location (38%, 57% and 44%). Consistent with these findings, the largest reduction in estimated AIS2+ injury probability, as calculated using the BrIC, was also observed at the pYrot location. Across helmet types, RLS-equipped helmets were associated with a probability of sustaining an AIS2+ injury after an impact at the pYrot location of 1-4%, corresponding to a Δ_*RLS*_ of 93-98% relative to conventional helmet configurations. It is worth noting that while pZrot is described as the least effective location for RLS technology, it still provides a Δ_*RLS*_ in p(AIS2+) of 47-78% across helmet types.

### 3.3 RLS effectiveness across headforms

The HIII headform was used for the majority of the testing presented in this study to enable direct comparison with existing literature ^24,34^. In line with the second objective of the study, additional impact testing of the urban helmet type was conducted using the EN 17950 headform to evaluate whether the RLS effectiveness across kinematics and injury metrics is sensitive to headform selection. Across both headforms, Δ_*RLS*_ in PLA was small, whereas an increased RLS effectiveness was observed for rotational kinematics (PAV and PAA) and the associated injury metrics (BrIC and p(AIS2+)) (Table 3, Figure 5). The highest RLS effectiveness across metrics was observed at the pYrot impact location for both headforms, consistently to what was observed in Section 3.2.

**Table 3.** Mean percentage reduction in kinematics and injury metrics between conventional and RLS-equipped urban helmets, Δ_*RLS*_ obtained by impact testing with two different headforms, HIII and EN 17950. Permutation tests were implemented to assess whether Δ_*RLS*_ is sensitive to headform selection. 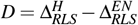 and its 95% CI were reported together with two-sided p-values. Statistical differences (*p <* 0.05) are shown in bold.

| Metric | pXrot |  | pYrot |  | pZrot |  |
| --- | --- | --- | --- | --- | --- | --- |
|  | Hybrid III | EN 17950 | Hybrid III | EN 17950 | Hybrid III | EN 17950 |
| $\Delta_{RLS}$ in PLA (%) | 4 ± 6<br>$D = 3\%[-20, 25], p = 0.796$ | 2 ± 4 | 8 ± 1<br>$D = -1\%[-17, 14], p = 0.821$ | 10 ± 4 | 1 ± 5<br>$D = -6\%[-14, 2], p = 0.161$ | 7 ± 4 |
| $\Delta_{RLS}$ in PAV (%) | 49 ± 4<br>$D = 2\%[-10, 14], p = 0.739$ | 46 ± 4 | 85 ± 4<br>$D = 5\%[0, 11], p = 0.057$ | 80 ± 3 | 37 ± 2<br>$D = -4\%[-19, 11], p = 0.599$ | 42 ± 4 |
| $\Delta_{RLS}$ in PAA (%) | 24 ± 14<br>$D = -8\%[-27, 11], p = 0.429$ | 32 ± 5 | 76 ± 1<br>$D = -4\%[-10, 3], p = 0.309$ | 80 ± 6 | 32 ± 7<br>$D = -5\%[-18, 8], p = 0.491$ | 37 ± 5 |
| $\Delta_{RLS}$ in BrIC (%) | 35 ± 3<br>$D = 6\%[-11, 23], p = 0.502$ | 29 ± 6 | 85 ± 4<br>**$D = 6\%[0, 11], p = 0.026$** | 80 ± 6 | 37 ± 2<br>$D = -5\%[-18, 8], p = 0.448$ | 42 ± 4 |
| $\Delta_{RLS}$ in p(AIS2+) (%) | 63 ± 4<br>$D = 5\%[-18, 28], p = 0.663$ | 58 ± 8 | 99 ± 1<br>$D = 1\%[0, 2], p = 0.052$ | 98 ± 1 | 47 ± 2<br>$D = -16\%[-32, 1], p = 0.063$ | 63 ± 6 |

**Fig. 5.**
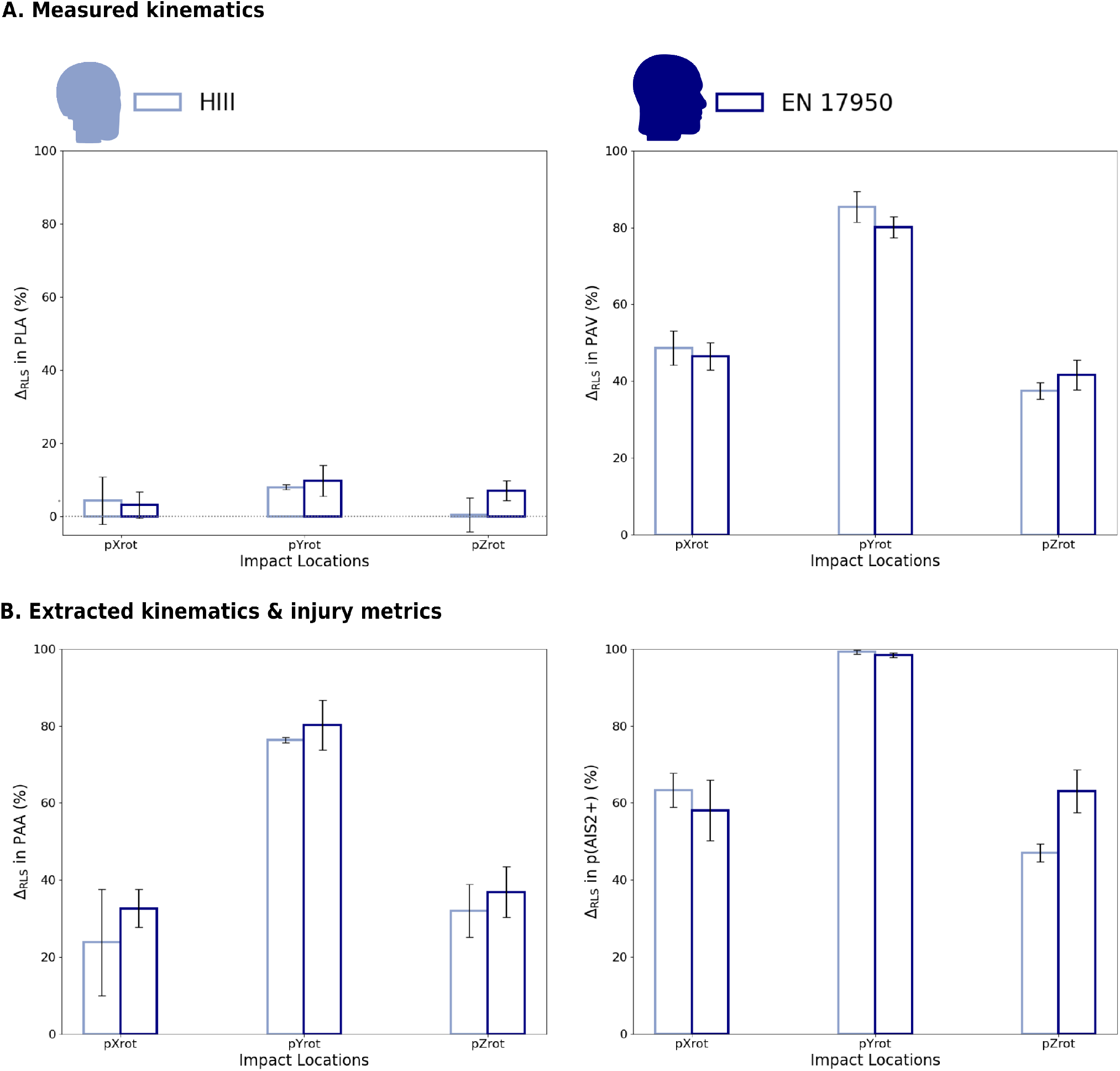
Mean percentage reduction by RLS (Δ_*RLS*_) for urban helmets, tested with the HIII (light blue) and the EN17950 (navy) headforms across all metrics and impact locations. Error bars indicate uncertainty in the estimated percentage reduction.

The difference in RLS effectiveness between headforms, *D*, varied by metric and impact location. In most cases, the magnitude of *D* was less than 10% and there was no consistent direction of the effect, as indicated by changes in its sign across metrics and locations. For the rotational kinematics metrics (PAV and PAA), permutation tests did not identify statistically significant differences at any impact location, and confidence intervals for *D* included zero in all cases. For the injury metrics, differences between headforms were likewise generally modest. Most met-ric–impact location combinations yielded non-significant permutation test results, with confidence intervals for *D* spanning zero, indicating insufficient evidence of sensitivity to headform selection. An exception was observed for BrIC at the pYrot location, where the estimated difference favored the HIII headform and the permutation test indicated a statistically significant difference (*D* = 6%, *p* = 0.026). Another notable observation was made for the p(AIS2+) metric at the pZrot location, where the estimated effect size, *D*, was comparatively large (16%) and visually apparent as the only case of non-overlapping error bars in Figure 5. The corresponding permutation test did not reach statistical significance and the confidence interval included zero. This result was therefore interpreted as suggestive but inconclusive. Overall, permutation testing indicated that, within the resolution of the present dataset, the RLS technology was largely headform-agnostic in its ability to mitigate kinematics and associated injury metrics. Statistically significant headform-dependent differences in percentage reduction were limited to specific metric–impact combinations, most notably BrIC in pYrot impacts, while other observed differences were accompanied by confidence intervals that included zero.

## 4 Discussion

This study evaluated the effectiveness of cycling helmets equipped with the Release Layer System (RLS) compared with conventional EPS helmets, with a focus on rotational kinematics and the associated risk of mTBI. Across three helmet types and multiple impact locations, RLS integration substantially reduced rotational metrics and the corresponding injury risk, while producing smaller and less consistent improvements in linear kinematics. The following discussion expands on these findings by examining their broader implications, contributing factors, and areas for further investigation.

### 4.1 Assessment of conventional helmet performance

The conventional helmets of all three helmet types tested well below the limit of the EN 1078:2012 standard in terms of PLA (250 g) ^17^, and their PAV and PAA results were consistent with those reported in previous studies of commercial helmets ^27–29^, suggesting that they are a representative baseline. The rotational performance of the conventional helmets corresponded to an average estimated AIS2+ injury probability of 64-68% across all impact locations, motivating the need for improved rotational protection. Integration of the RLS technology reduced rotational kinematics (Δ_*RLS*_ of 57-66% in PAV and 44-56% in PAA) across all helmet types, yielding a corresponding reduction in AIS2+ injury probability to 9–22% (Δ_*RLS*_ of 68–86%). Reductions in PLA (4.5–9.7%) were also observed, although statistical significance was reached only for the road helmet. Both this observation and the variability seen between impact locations (black circles, triangles, and squares representing pXrot, pYrot, and pZrot, respec-tively, Figure 3) highlight the need for further investigation to better understand the influence of helmet geometry and RLS integration on impact location-specific performance.

### 4.2 Repeatability of test results

Assessing the results at the level of individual impact locations allows for evaluation of repeatability across helmet types and configurations. The coefficient of variation (CV) for kinematic and injury metrics was found to be below 10% for the majority of tests (100%, 78%, 83%, 83%, and 61% of helmets for PLA, PAV, PAA, BrIC, and p(AIS2+), respectively; Figure S2A in Supplementary Information). Instances where CV exceeded 10% were limited to RLS-equipped helmets, with higher variability observed in PAV and p(AIS2+) at the pYrot and pZrot locations. This increase might be expected for pYrot, as the low mean values of rotational kinematics at this location elevate CV even when standard deviations remain low (CV = standard deviation*/*mean). Examining standard deviations provides a more direct comparison of the variability in our results with intralaboratory variability reported in literature. In our dataset, standard deviations reached up to 2.6 rad/s for PAV, 850 rad*/*s^2^ for PAA, and 0.08 for p(AIS2+), values consistent with those in the literature ^27,31,46^(Figure S2B in Supplementary Information). The largest standard deviation for PAV, and thus p(AIS2+), was observed in the pZrot location of the road helmet, likely due to the complex dynamics of this test configuration. Excluding this case, standard deviations for PAV reached up to 1.6 rad/s. Overall, the results demonstrate repeatability across impact locations, with the observed variability being consistent with published intralaboratory ranges. This can further be observed in the time-series data of the resultant head kinematics (Figures S3 and S4 in Supplementary Information).

### 4.3 Performance across impact locations

Accident databases and oblique impact testing have shown that impact location plays an important role in brain injury prevalence and severity ^22,30,47,48^. In our dataset, impact-location-dependent differences in rotational kinematics were observed across helmet types (Figure 4). For RLS-equipped helmets, reductions in rotational kinematics were generally larger at the pYrot location, then the pXrot location, and finally the pZrot location, whereas conventional helmets exhibited different relative responses across locations. These observations were consistent across both headforms tested (Table 3, Figure 5, Table S3, Figure S1), indicating that impact location and helmet–system interactions both influence performance.

While no formal statistical analysis of interaction effects between impact location and helmet configuration was performed, due to the limited number of observations per condition, the observed location-dependent differences are consistent with prior studies on other rotation-mitigation technologies ^29,30^. Bonin et al. ^30^ hypothesised that greater reductions at the pYrot location are linked to the larger radius of curvature of the headform and helmet in the sagittal plane, resulting in a longer arc length for rotations about the y-axis compared with the x-axis. In the context of RLS, this hypothesis is qualitatively supported by the orientation of the panels, which are aligned with the helmet’s longitudinal axis. During a pYrot impact, the panel releases along this axis and follows the larger radius of the headform, remaining in contact with the anvil-headform interface. This configuration allows more spheres to deploy and dissipate energy through rolling. In contrast, during a pXrot impact, the shorter arc length and release along the helmet’s shorter axis provide less coverage of the interface, limiting the number of spheres deployed and the energy dissipated. Integration details can modify this behaviour, as multiple panels may be released depending on the helmet type.

The mechanism differs for pZrot impacts, which involve compound rotations about both the z- and y-axes. The complex kinematics and reduced panel coverage at the impact site likely contribute to the lower effectiveness of RLS in this orientation, though Δ_*RLS*_ of 37–58% in PAV was still achieved. In addition, rotation about the y-axis shifts the headform’s center of gravity out of balance relative to the helmet, potentially increasing sensitivity to positioning prior to impact testing.

### 4.4 Influence of helmet geometry on RLS effectiveness

The RLS mechanism of operation might be partly responsible for the trends in RLS effectiveness across impact locations and irrespective of helmet type described above, however differences in helmet geometry also appear to influence performance. In particular, the large, uniform A-surface of the urban helmet, together with its curvature and lack of vents at the pYrot location, provides favourable conditions for RLS deployment, reflected in a 98% reduction in the risk of sustaining an AIS2+ injury at this location (Table 2). The road and MTB helmets, on the other hand, feature narrower panels at pYrot to accommodate ventilation, reducing RLS coverage and increasing residual kinematics relative to the urban helmet, though rotation-mitigation remains large (97% and 93% reduction in the risk of sustaining an AIS2+ injury, respectively, Table 2). At the pXrot location, the road and MTB helmets again share similar geometric features—narrow RLS panels combined with vent openings—resulting in comparable rotational performance. These observations demonstrate that RLS deployment and thus effectiveness is shaped by the interaction of impact location and helmet geometry effects. Tuning of the integration of the technology can therefore potentially optimize performance within the design constraints of each helmet type.

### 4.5 RLS integration effects on linear kinematics

Unlike the reductions in PAV, PLA attenuation did not show any consistency between helmet types. In 3.1, a significant Δ_*RLS*_ in PLA was detected for road helmets, and in 3.2 and Figure 4, significant reductions were identified across multiple impact locations for all helmet types. This indicates that the influence of RLS on peak linear kinematics is likely more complex. Various hypotheses can be put forward to explain the underlying mechanism. One possibility is that the additional shell functions similarly to the internal injection-molded reinforcements found in low-profile road helmets, stiffening the EPS and limiting splaying under load. The placement of RLS panels may then determine where this reinforcement is most effective. Another hypothesis is that localized forces at the contact points of the polycarbonate spheres induce small-scale surface rippling of the underlying EPS, leading to locally increased compression and densification.

This effect is analogous to micro-scale coring, in which discrete regions of EPS undergo enhanced deformation without material removal. Further experimental work is needed to clarify these mechanisms and to establish whether the observed reductions in PLA represent a consistent and reproducible benefit of RLS integration across helmet types and impact conditions.

### 4.6 Headform influence

The results in 3.3 indicate that Δ_*RLS*_ in PLA, PAV and PAA was not affected by the headform used. This outcome was expected, as differences in moment of inertia (MoI) and coefficient of friction (CoF) between headforms were hypothesised to influence conventional and RLS-equipped helmets in a similar manner, given that RLS is a thin, outer layer and does not directly interact with the headform surface. Previous studies have shown that MoI and CoF can affect rotational kinematics ^30,35,37,49–51^, though their relative importance remains debated. Yu et al. ^51^ further reported a significant interaction between headform and impact location for all injury metrics except brain strain, highlighting the complexity of these factors. Further work is needed to: (a) clarify the relative contributions of MoI and CoF, and (b) determine how CoF may vary over time and orientation, to ensure that the EN 17950 remains biofidelic not only after cleaning but also across repeated testing. This requirement is reflected in the EN 17950 protocol, which specifies CoF measurement along both long and short axes ^38^.

Although headform choice did not significantly affect most peak kinematics and injury metrics (p-values *>* 0.05 for most locations and zero included in the 95% CIs of *D*), permutation tests did find a significant difference in BrIC at pYrot. Moreover, in three instances (PAV and p(AIS2+) at pYrot, and p(AIS2+) at pZrot), the p-values were close to the significance threshold (0.057, 0.052, and 0.063, respectively), suggesting that further investigation with a larger sample size is warranted to confirm these findings. Such findings also emphasize the need for future studies to assess the influence of headform selection on brain injury risk estimation using multiple injury metrics or computational brain models ^52,53^. Detailed analysis of the kinematic time-series data, rather than just peak values, using tools such as ARC-Gen ^54^, may also provide further insights into the biomechanical implications of headform selection. As a preliminary observation, differences were noted in the shape of certain kinematic traces obtained with different headforms (Figures S3 and S4 in Supplementary Information).

### 4.7 Limitations and future work

The scope of this study was subject to certain limitations, which should be acknowledged alongside the opportunities they present for future work. First, all tests were conducted at an impact speed of 6.5 m/s, which is consistent with standard helmet testing. Additional testing at higher and lower speeds would provide a more complete understanding of RLS performance under different cycling accident scenarios reported in literature ^42,48,55,56^. Second, impact tests were conducted at the pXrot, pYrot, and pZrot locations, representing the front and side regions of the helmet most commonly impacted ^22^. Future work should also include impact testing at the rear of the helmet, reported in the literature as a relevant area for cyclist head injuries ^22^, and evaluate whether the nYrot impact location (symmetric to pYrot) appropriately represents accidents affecting the rear of the head ^55^.

The comparative headform analysis also has constraints. Data was collected only for the urban helmet, selected because it exhibited the smallest reduction in rotational kinematics averaged across locations (Table 1) and the largest variation in RLS effectiveness between locations (Table 2). While this allowed assessment of both the upper and lower bounds of RLS effectiveness, extending the analysis to include road and MTB helmets would provide a broader evaluation. Moreover, the study assessed the combined influence of headforms and their instrumentation, as different sensor arrays were used. All sensors were calibrated and the same data filtering and analysis protocols were applied, so significant variations due to the sensor arrays are not expected. Another limitation arises from testing across different laboratories. Experiments with the HIII headform were performed at UNISTRA, whereas EN 17950 testing was conducted at HEXR. Although identical protocols were followed, collecting additional EN 17950 data at UNISTRA would have enabled a more direct comparison. This was not feasible due to limited availability of EN 17950 headforms and compatible instrumentation at the time of the study. The interaction between headform characteristics and impact location also warrants further study. Yu et al. ^51^ reported significant interactions using an earlier version of the EN 17950, produced by a different manufacturer and tested under earlier validation procedures. Extending our analysis to validate such statistical models with the current, commercially available EN 17950 would be valuable.

Finally, the study focuses on differences in rotational kinematics between conventional and RLS-equipped helmets across helmet types and headforms. To further contextualise our findings, we calculated BrIC, a metric widely used in literature, and used it to estimate the probability of AIS2+ injuries, allowing comparison with other studies evaluating the effectiveness of helmets with rotation-mitigation technologies. While BrIC is commonly applied in the cycling head protection field ^27–31^, we acknowledge that its mapping to injury risk relies on car accident data, which may limit its direct applicability to cycling head injuries. As a result, future work should investigate how the effectiveness of RLS in reducing AIS2+ risk varies when alternative injury and tissue-level metrics are employed ^52,53,57^.

## 5 Conclusions

The present study evaluated the effectiveness of the Release Layer System (RLS), a novel rotation-mitigation technology integrated onto urban, road and MTB helmets, compared with conventional EPS helmets of equivalent geometry. Oblique impact testing was conducted across three helmet types, three impact locations, and two headforms (n = 96). RLS-equipped helmets exhibited peak linear accelerations similar to EPS models, with only minor improvements in some cases. In contrast, RLS consistently reduced peak angular velocity, with average reductions of 57–66% across helmet types and locations, corresponding to a 68–86% decrease in the risk of AIS2+ injury. The magnitude of this reduction varied by impact location, with the largest improvements observed in pYrot. Comparative testing with the HIII and the new EN 17950 headforms for one helmet type showed no systematic differences in percentage reductions of angular metrics, though some location-specific differences in injury risk were noted. These findings highlight the potential of RLS technology to substantially reduce rotational injury risk, while also underscoring the need for further work to interpret headform-specific outcomes.

## Acknowledgements

The authors would like to acknowledge Nicolas Bourdet, Caroline Deck and Rémy Willinger of the iCUBE laboratory of the University of Stras-bound (UNISTRA) for the collection of the oblique impact testing data and for valuable discussions of the results. The authors would also like to thank the entire HEXR team for their contributions to the development of the RLS technology, insightful comments, and assistance throughout the study; in particular: James Humphries and Darcy Thompson-Bagshaw. Finally, the authors would like to thank Canyon Bicycles for their support in supplying the MTB helmets.

