## Supplementary Information for "A Novel Rotation-Mitigation Technology for Cycling Helmets Tested Across Helmet Types, Impact Locations and Headforms"

#### 1 Brain Injury Criterion (BrIC) and probability of $AIS2^+$ injury[1]

$$BrIC = \sqrt{\left(\frac{\omega_x}{\omega_{xC}}\right)^2 + \left(\frac{\omega_y}{\omega_{yC}}\right)^2 + \left(\frac{\omega_z}{\omega_{zC}}\right)^2}, \tag{1}$$

where $\omega_{xC} = 66.25$, $\omega_{yC} = 56.45$ and $\omega_{zC} = 42.87$.

$$p(AIS2+) = 1 - e^{-\left(\frac{BrIC}{0.602}\right)^{2.84}} \tag{2}$$

#### 2 Materials & Methods

Table S1 Physical characteristics of the helmet types and configurations tested in this study and an overview of the headforms used for testing.

| Helmet | Weight (g) | Weight of RLS technology (g) | Additional weight (g) | Headform used for testing |
|---|---|---|---|---|
| Urban | 300 ±5 | - | - | HIII & EN 17950 |
| Urban-RLS | 345 ±3 | 15 | 30 | HIII & EN 17950 |
| Road | 263 ±5 | - | - | HIII |
| Road-RLS | 306 ±14 | 15 | 28 | HIII |
| MTB | 265 ±3 | - | - | HIII |
| MTB-RLS | 317 ±4 | 13 | 39 | HIII |

Table S2 Coefficient of friction (CoF) measurements for the EN 17950 headform before and after impact testing[2]. One-sample z-tests show no statistical differences of the CoF before impact testing from the prescribed CoF of $0.30 \pm 0.03$ ($Z = 1.04$, $n = 5$, $p = 0.297$ and $Z = 0.60$, $n = 5$, $p = 0.551$ for the right-to-left and back-to-front test orientations, respectively). One-way Welch ANOVAs also show no statistical difference of the CoF before and after impact testing.

| | Right-to-left | | Back-to-front | |
|---|---|---|---|---|
| Test cycle | Before impact testing | After impact testing | Before impact testing | After impact testing |
| 1 | 0.30 | 0.29 | 0.28 | 0.33 |
| 2 | 0.34 | 0.27 | 0.31 | 0.31 |
| 3 | 0.31 | 0.33 | 0.33 | 0.32 |
| 4 | 0.29 | 0.27 | 0.33 | 0.31 |
| 5 | 0.33 | 0.30 | 0.29 | 0.32 |
| **Test sequence mean** | $0.31 \pm 0.02$ | $0.29 \pm 0.02$ | $0.31 \pm 0.02$ | $0.32 \pm 0.01$ |
| | $F(1, 7.75) = 2.30$, $p = 0.169$ | | $F(1, 5.06) = 0.85$, $p = 0.400$ | |

#### 3 Results

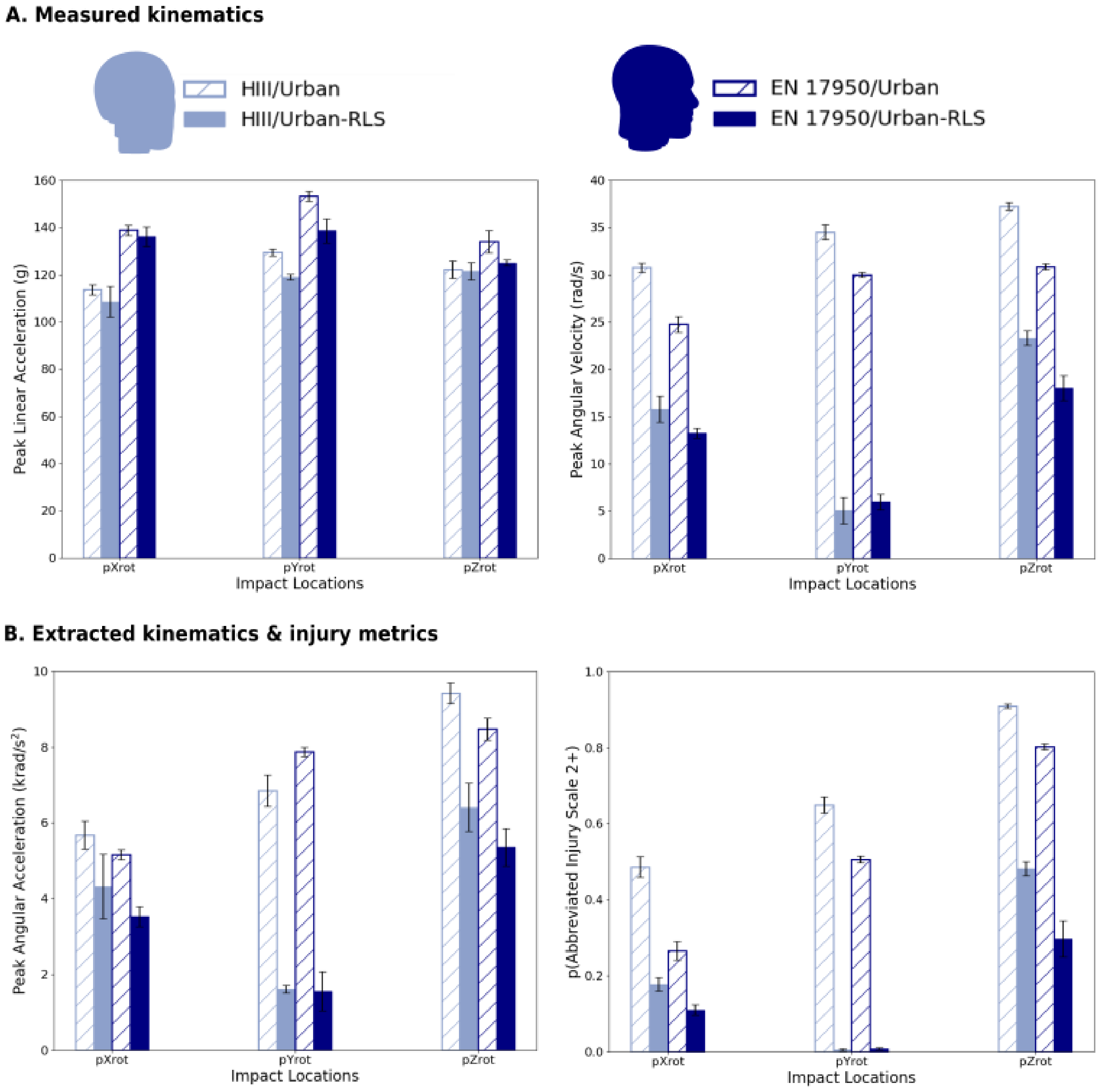


Fig. S1 Peak kinematics and injury metrics of conventional (hatched) and RLS-equipped (solid) urban helmets, tested with the HIII headform (light blue) and the EN17950 headform (navy).

Table S3 Mean and standard deviation values for conventional urban helmets (Control), RLS-equipped urban helmets (RLS), and mean percentage reduction between conventional and RLS-equipped urban helmets ($\Delta_{RLS}$) collected from impact testing with two different headforms, HIII and EN 17950. A permutation test was implemented to assess whether $D = \Delta^{H}_{RLS} - \Delta^{EN}_{RLS}$ is significant at the $\alpha = 0.05$ level (4900 exact permutations evaluated). $D$, 95% CI and two-sided p-values reported; statistically significant values shown in bold.

| | | pXrot | | pYrot | | pZrot | |
|---|---|---|---|---|---|---|---|
| Metric | | Hybrid III | EN 17950 | Hybrid III | EN 17950 | Hybrid III | EN 17950 |
| PLA | Control (g) | 113.7±2.1 | 138.9±2.1 | 129.5±1.5 | 153.3±2.1 | 122.2±3.7 | 134.0±4.7 |
| | RLS (g) | 108.5±6.6 | 136.2±4.0 | 119.0±1.1 | 138.6±5.1 | 121.5±3.6 | 125.0±1.3 |
| | $\Delta_{RLS}$(%) | 4±6 | 2±4 | 8±1 | 10±4 | 1±5 | 7±4 |
| | | $D = 3\%[-20, 25]$, $p = 0.796$ | | $D = -1\%[-17, 14]$, $p = 0.821$ | | $D = -6\%[-14, 2]$, $p = 0.161$ | |
| PAV | Control (rad/s) | 30.8±0.5 | 24.8±0.8 | 34.5±0.8 | 30.0±0.3 | 37.3±0.4 | 30.9±0.3 |
| | RLS (rad/s) | 15.8±1.4 | 13.2±0.6 | 5.1±1.4 | 6.0±0.8 | 23.3±0.8 | 18.0±1.3 |
| | $\Delta_{RLS}$(%) | 49±4 | 46±4 | 85±4 | 80±3 | 37±2 | 42±4 |
| | | $D = 2\%[-10, 14]$, $p = 0.739$ | | $D = 5\%[0, 11]$, $p = 0.057$ | | $D = -4\%[-19, 11]$, $p = 0.599$ | |
| PAA | Control (krad/s$^2$) | 5.67±0.37 | 5.17±0.13 | 6.85±0.42 | 7.88±0.12 | 9.43±0.27 | 8.48±0.29 |
| | RLS (krad/s$^2$) | 4.33±0.85 | 3.52±0.26 | 1.61±0.10 | 1.56±0.51 | 6.41±0.64 | 5.36±0.50 |
| | $\Delta_{RLS}$(%) | 24±14 | 32±5 | 76±1 | 80±6 | 32±7 | 37±5 |
| | | $D = -8\%[-27, 11]$, $p = 0.429$ | | $D = -4\%[-10, 3]$, $p = 0.309$ | | $D = -5\%[-18, 8]$, $p = 0.491$ | |
| BrIC | Control | 0.52±0.01 | 0.40±0.01 | 0.61±0.01 | 0.53±0.00 | 0.82±0.01 | 0.71±0.01 |
| | RLS | 0.34±0.01 | 0.28±0.01 | 0.09±0.02 | 0.11±0.02 | 0.52±0.01 | 0.42±0.03 |
| | $\Delta_{RLS}$(%) | 35±3 | 29±6 | **85±4** | **80±6** | 37±2 | 42±4 |
| | | $D = 6\%[-11, 23]$, $p = 0.502$ | | $\mathbf{D = 6\%[0, 11], p = 0.026}$ | | $D = -5\%[-18, 8]$, $p = 0.448$ | |
| p(AIS2+) | Control | 0.49±0.03 | 0.27±0.02 | 0.65±0.02 | 0.51±0.01 | 0.91±0.01 | 0.80±0.01 |
| | RLS | 0.18±0.02 | 0.11±0.01 | 0.01±0.00 | 0.01±0.00 | 0.48±0.02 | 0.30±0.05 |
| | $\Delta_{RLS}$(%) | 63±4 | 58±8 | 99±1 | 98±1 | 47±2 | 63±6 |
| | | $D = 5\%[-18, 28]$, $p = 0.663$ | | $D = 1\%[0, 2]$, $p = 0.052$ | | $D = -16\%[-32, 1]$, $p = 0.063$ | |

### 4 Discussion

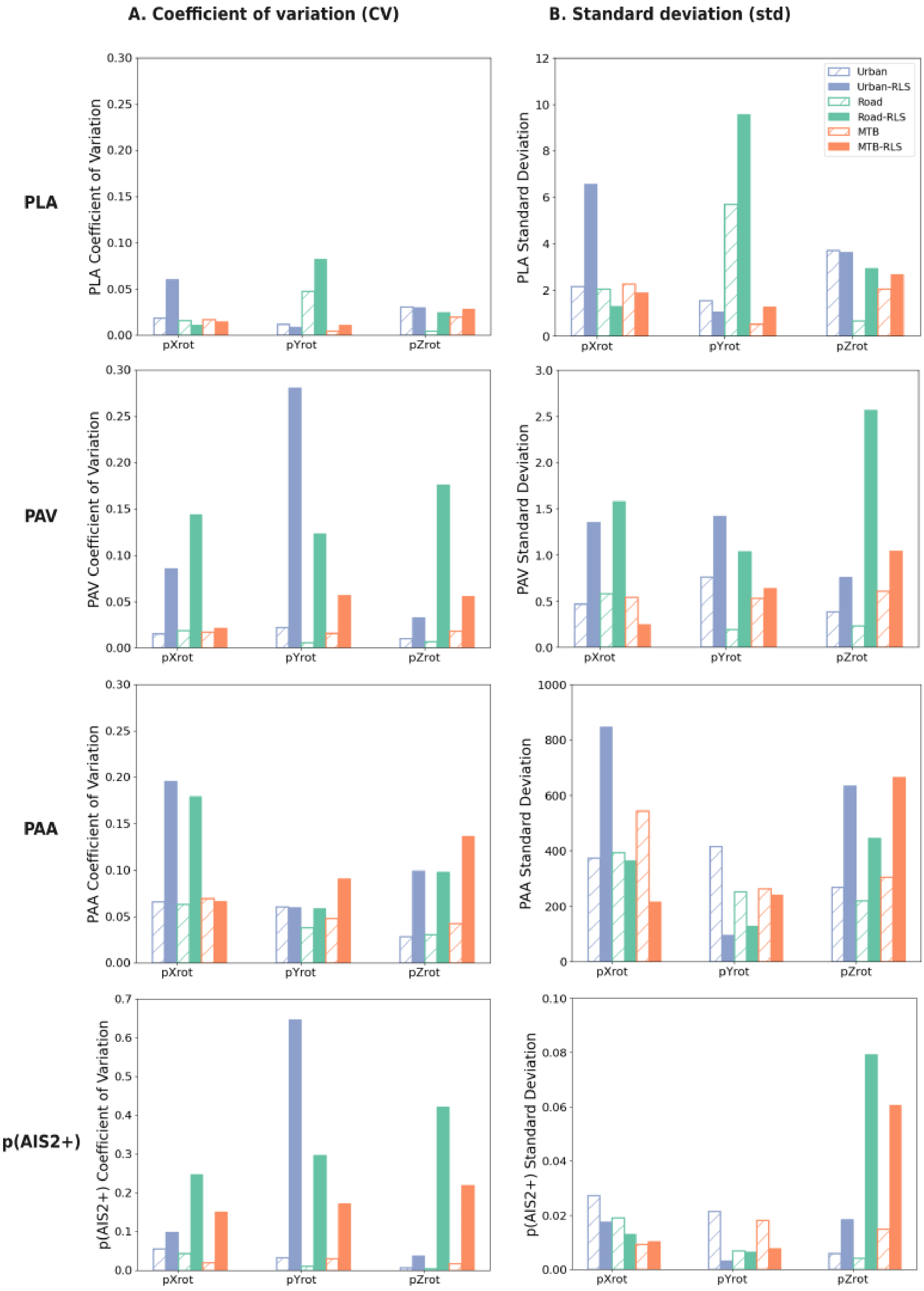


Fig. S2 Coefficient of variation (CV) and standard deviation (std) by impact location for three helmet types across measured kinematics, and extracted kinematics and injury metrics.

#### Kinematic traces per impact location - Hybrid III

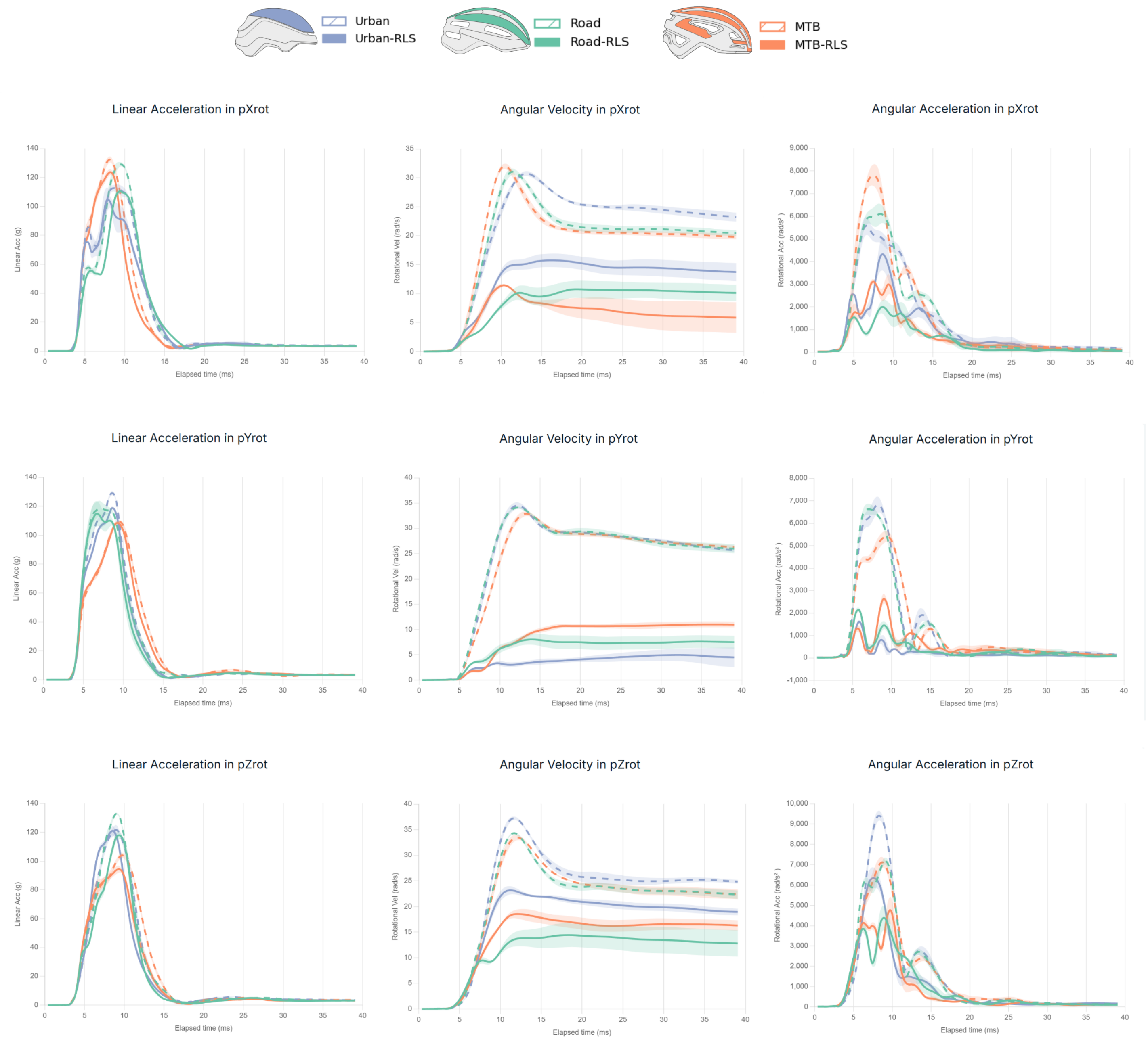


Fig. S3 Mean ± 95% CI of head kinematics (resultant linear acceleration, angular velocity and angular acceleration) for all impact locations and helmet types tested using the HIII headform.

#### Kinematic traces per impact location - EN 17950

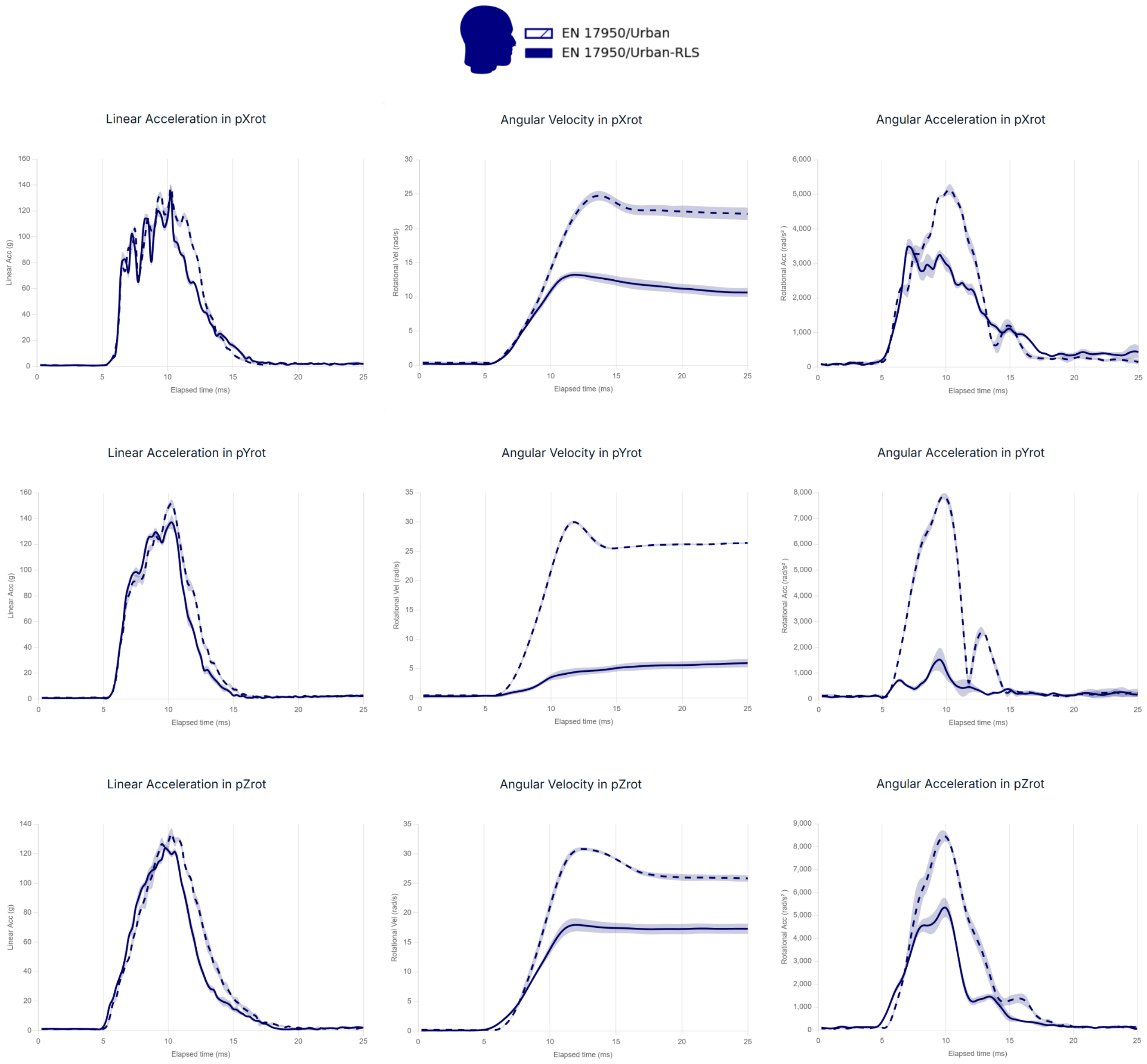


Fig. S4 Mean $\pm 95\%$ CI of head kinematics (resultant linear acceleration, angular velocity and angular acceleration) for all impact locations for the urban helmet type tested using the EN 17950 headform.

[2] European Committee for Standardization. EN 17950: Protective helmets — Test methods — Shock absorption including measuring rotational kinematics. Brussels, Belgium: European Committee for Standardization; 2024. EN 17950.
